# Clear Cell Renal Cell Carcinoma with Biallelic Inactivation of CDKN2A/B on 9p21 have Distinct Gene Expression Signature and are Associated with Poor Prognosis

**DOI:** 10.1101/143180

**Authors:** Andrew H. Girgis

## Abstract

**Purpose:** The clinical implications of biallelic inactivation of *CDKN2A/B* in clear cell renal cell carcinoma (ccRCC) and relevant dysregulated biological pathways and gene signatures were investigated.

**Materials and Methods:** Data were obtained from the TCGA data set and validated using Project GENIE and previously published dataset. *CDKN2A/B* allelic status was classified into 3 groups, including biallelic *CDKN2A/B* inactivation (homozygous deletion or combined heterozygous deletion and mutation), monoallelic *CDKN2A/B* loss (heterozygous deletion or mutation) and absent *CDKN2A/B* allelic loss. Univariate and multivariate cancer-specific survival and disease-free survival analyses were performed. Integrated analyses of copy number, gene expression (mRNA and miRNA), protein expression and methylation changes were conducted.

**Results:** Of 440 patients with ccRCC 17 (3.9%) had biallelic *CDKN2A/B* inactivation and 116 (26.4%) had monoallelic *CDKN2A/B* loss. *CDKN2A/B* allelic inactivation was associated with late tumor stage, high histological grade, presence of metastasis and greater tumor size. Patients with biallelic deletion of *CDKN2A/B* showed significantly worse cancer-specific survival and disease-free survival (p<0.0001). Significant co-occurrence of *MTAP* homozygous deletion was observed in *CDKN2A/B* biallic inactivated tumors (46.7%; p<0.001). Significant underexpression of *CDKN2A/B* was observed in biallelic inactivated tumors at the mRNA and protein levels. miR-21 was the most highly expressed miRNA in biallelic inactivated tumors. Biallelic inactivated tumors were significantly enriched for genes related to activation of *ATR* in response to replication stress and miR-21 target genes.

**Conclusions:** *CDKN2A/B* biallelic inactivation may be a prognostic marker for ccRCC and is associated with distinct dysregulation of gene expression signatures.

## INTRODUCTION

Renal cell carcinoma (RCC) is the most prevalent neoplasm of the adult kidney and the incidence of RCC continues to increase (1–3). Alarmingly, and despite advances in early detection and treatment, the incidence of RCC is not accompanied by a decrease in the rate of mortality, (3). RCC is comprised of several histopathological subtypes, of which the most common (~75%) is the clear cell subtype (ccRCC), where the biallelic inactivation of *VHL* presents in most sporadic ccRCC cases. However, in the absence of other markers, the limitations of histology for this subtype define a challenge hindering clinicians from gauging appropriately the clinical outcome and treatment response, as ccRCC is remarkably heterogeneous at the molecular level. Furthermore, the coordinated interplay of multiple regulatory mechanisms (copy-number alterations, methylation changes, mutations, miRNA regulation, etc.) on cancer-associated genes is also poorly understood in these cancers. Thus, determining the influence of genetic and epigenetic alterations on the expression of cancer related genes and their impact on biologic pathways remains a complex endeavor in ccRCC. Deeper delineation of the molecular pathology of ccRCC by surveying and investigating these genetic and epigenetic changes and their effects on key molecules and their respective biologic pathways is of crucial importance for the improvement of current diagnostics, prognostics, and the tailoring of treatment of options (4).

Recent large-scale efforts from The Cancer Genome Atlas (TCGA) cataloguing 400 ccRCC tumors for genetic and epigenetic alterations have revealed novel mutations of *SETD2, BAP1, ARID1A* and *PBRM1*, among other significantly mutated genes with less prevalence than *VHL* (5). Notably, mutations of the tumor suppressors, *BAP1* and *SETD2*, located at 3p21, were associated with adverse clinical outcomes in ccRCC. This locus is subject to copy-number loss in the majority of ccRCC cases and, thus, complete inactivation by combination of mutation and copy-number loss of these tumor suppressors has proved to be prevalent (6). Furthermore, a study using the TCGA dataset demonstrated that the tumor suppressor *PTEN* is subject to biallelic inactivation (combined copy number loss and mutation) in 2.6% of cases. This scenario was significantly associated with worse overall survival and higher stage and grade compared to tumors with monoallelic loss and diploid tumors (7). Hence, identifying tumor suppressors subject to biallelic inactivation in ccRCC may provide insight into clinically and biologically relevant subpopulations of tumors to help explain their observed clinical variability.

The 9p21 region is implicated in a variety of cancers and encodes for the tumor suppressor genes *CDKN2A/2B* and their associated isoforms: p16INK4a, p14ARF, and p15INK4b *(CDKN2B)*. These proteins regulate progression of cellular proliferation through blockade of cyclin/CDK-4/6 complexes, preventing cell division during the G1/S phase of the cell cycle and perturbation of their synthesis may lead to dysregulation of cell growth (8,9). Several studies, including the TCGA, have identified homozygous deletion of the tumor suppressors *CDKN2A* and *CDKN2B* in ccRCC (10–12). Loss of *CDKN2A* in ccRCC is due in part to monoallelic deletion (loss of heterozygosity, LOH) and is associated with poor prognosis, increased stage and metastasis (13,14). Furthermore, a recent study reported 9p monoallelic loss as an independent marker of risk of recurrence and RCC-related mortality (15). However, complete inactivation of *CDKN2A* is relatively rare in ccRCC as *CDKN2A* mutation and homozygous deletion is uncommon (5). Also, more recently, loss-of-function germline mutations of *CDKN2B* were shown to predispose to RCC providing further evidence of the importance of biallelic inactivation of *CDKN2B* as a driver to carcinogenesis of RCC (16). A detailed constitutive analysis of *CDKN2A/B* biallelic inactivation in association with genome-wide genetic and epigenetic alterations, as well as clinical data was performed, to determine the importance of *CDKN2A/B* inactivation in ccRCC.

## MATERIALS AND METHODS

### TCGA Data Acquisition

Data were obtained from level 3 TCGA data sets from the ‘KIRC Archives’, including SNP copy-number, gene and miRNA expression, methylation, and protein and clinical data using the January 2016 analysis run firehose from the Broad Institute’s Genome Data Analysis Centre (http://gdac.broadinstitute.org/runs/stddata_2016_01_28). TCGA data types, platforms, and methodologies are as described previously (5). Briefly, data were generated at TCGA Genome Data Analysis Centers. Whole exome sequencing was performed with the Illumina^®^ and SOLiD^®^ platforms. Putative copy number alterations were estimated using the GISTIC2.0 method with the Affymetrix^®^ SNP 6 platform (17). Normalized RSEM was used to estimate mRNA expression (18). RNAseq data were generated with the IlluminaGA_RNASeqV2 or IlluminaHiSeq_RNASeqV2 platform. Protein expression was estimated at the M.D. Anderson Reverse Phase Protein Array Core Facility. A subset of TCGA cases containing recently misdiagnosed ccRCCs (identified as chromophobe, clear cell papillary and translocation RCCs (19,20)) was removed from analysis. The data were limited to the number of cases previously analyzed by the TCGA Research Network for a total of 440 cases (19–21).

### Validation sets from Project GENIE and Sato *et al.*

The ccRCC dataset of Project GENIE was analyzed for verification of findings as made available through the cBio Cancer Genomics Portal (http://cbioportal.org/genie/) (22–24). The Project GENIE dataset is comprised of tumors with data on sequenced mutations, high-resolution copy number, and clinical data including patient age, gender, race and ethnicity (n=237).

Copy number data of Affymetrix 250K SNP arrays were downloaded for 240 clear cell RCC samples from the Sato et al. dataset as previously published (https://www.ebi.ac.uk/arrayexpress/experiments/E-MTAB-2015/) (25). Clinical data was obtained from the Supplementary Table 1 of their publication. Normalization and segmentation were conducted using the Agilent GeneSpring GX 11 software with default settings.

### Subgrouping and analysis

Allelic status of *CDKN2A/B* in ccRCC tumors was classified as biallelic inactivation (homozygous deletion or combined monoallelic deletion and mutation), monoallelic loss (heterozygous deletion or mutation), or normal diploid (absence of allelic deletions and mutations). Possible associations of *CDKN2A/B* allelic status with tumor characteristics, as well as participant age, gender, ethnicity and race was assesssed. Assessment of statistical significance of age was estimated using ANOVA. The Fisher exact test was used for categorical variables, and the Kruskal-Wallis test for ordered variables. Cancer-specific survival and disease-free survival were compared among *CDKN2A/B* allelic status groups with the cumulative incidence of competing events and Gray Test. The multivariate Cox proportional hazards model was applied, adjusting for age, to AJCC tumor stage, grade and tumor size. All statistical analyses were conducted using XLSTAT (Addinsoft XLSTAT ver. 18.07; www.xlstat.com).

### Gene Set Association Analysis (GSAA) for RNASeq analysis

Gene Set Association Analysis (GSAA) was conducted to assess the distribution of underexpressed and overexpressed sets of genes in relation to their genomic location and identify effects of copy number losses or gains on mRNA expression for the three separate allelic phenotypes of *CDKN2A/B*: Biallelic inactivated, monoallelic lost and diploid tumors (26). Enrichment of gene sets was based on the Molecular Signatures Database version 5.1 (msigdb_v5.1.xml). GSAA was conducted between the allelic phenotypes with 1000 permutations, minimum priori of 5 genes, and nominal p value < 0.01. GSAA was also conducted for pathway and gene signature analysis to identify significantly enriched biological pathways and signatures from the literature.

### miRNA expression analysis

The “Level 3” TCGA miRNA dataset was used for analyzing differential expression in a total of 173 cases of the three different subgroups; biallelic inactivation, monoallelic loss and diploid tumors. Upper quartile normalization was applied on read count values as previously described to account for artefacts generated by large expression changes in the most prevalent miRNAs (27). Volcano plots were generated applying 0.5 (x) and 0.05 (y) thresholds on log2(Mean ratio) values and 5% significance level using XLSTAT (Addinsoft XLSTAT ver. 18.07; www.xlstat.com).

### Gene-specific Methylation analysis

Methylation changes for candidate genes were assessed using matched “Level 3” TCGA ccRCC methylation data. A b-value difference of 0.35 between tumor and normal kidney was selected as the threshold for hypermethylation (>0.35) or hypomethylation (<-0.35) as previously described (28). TCGA data types, platforms, and methodologies are as described previously (The Cancer Genome Atlas Research Network 2008).

## RESULTS

### Biallelic and Monoallelic Deletions of *CDKN2A/2B* in ccRCC

Fifteen cases (3.4%) of the 440 cases surveyed presented with homozygous deletion of *CDKN2A/B*, and 2 cases (0.5%) with a combined copy-number loss and deleterious mutation of *CDKN2A* (Figure 1). A total of 5 *CDKN2A* mutations in 5 of 440 ccRCC cases were observed (Supplementary Table 1). Missense *CDKN2A* mutations with at least a medium functional impact score or significant CHASM score (See Materials and Methods) were assumed to induce significant loss of function (29–31). Two cases presented with a missense mutation demonstrating a significant functional impact score. The remaining two deleteriously mutated cases coinciding with copy-number loss of *CDKN2A* presented with an in-frame insertion and nonsense mutation. No mutations were observed for *CDKN2B*. Monoallelic deletions of *CDKN2A/2B* were observed in 116 cases (26%). Low level gains were observed in 7 cases (1.6%). Further investigation of whether *CDKN2A/B* allelic status was associated with significant enrichment for regions of copy number alteration and mutations in ccRCCs was conducted. Genome-wide copy number changes and mutations in 440 ccRCCs from the TCGA (see Materials and Methods) were analyzed. Significant co-occurrence of *MTAP* homozygous deletion was observed with *CDKN2A* homozygous deletion (46.7%; p<0.001; Figure 1) consistent with previous reports of co-occurring deletion of these two genes (32).

**Figure 1.**
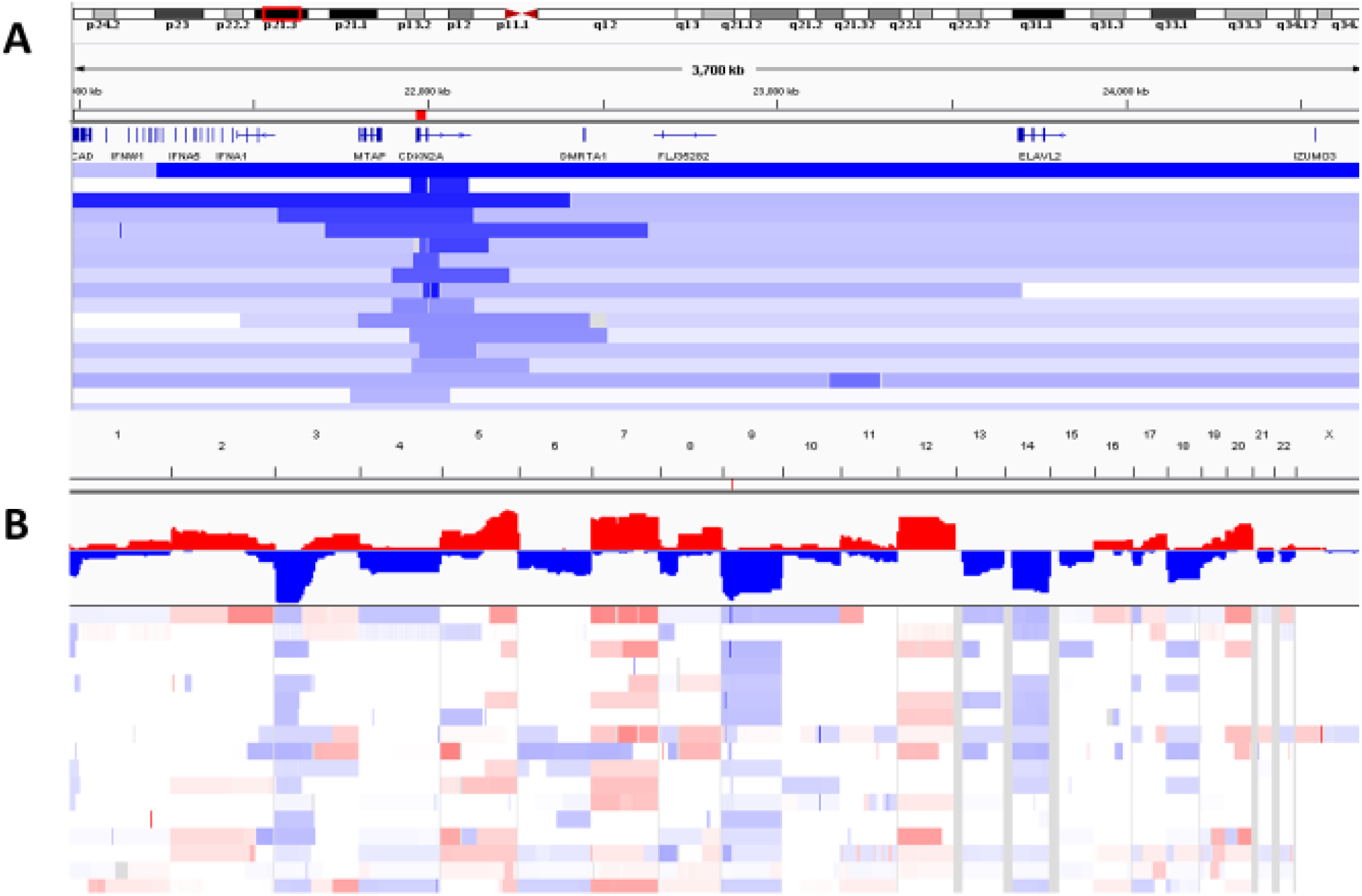
Biallelic deletions of CDKN2A/B in ccRCC from the TCGA Dataset. A, CDKN2A/B biallelic deletions mapped to genomic coordinates on 9p21 (deletions, blue). B, Copy number profiling of CDKN2A/B deleted ccRCC. Heatmap and frequency plot of CNAs as visualized by the Integrative Genomics Viewer; gains are in red and losses are in blue.

### Clinical Associations of *CDKN2A/2B* Allelic Status

A higher proportion of cases with monoallelic deletion were male (p <0.001), however no significant difference was observed for biallellic deletion. Allelic status was also significantly associated with age with higher median age among cases with biallelic deletion compared to diploid tumors (p = 0.047). Biallelic and monoallelic deletions were also significantly more frequent in higher grade and stage tumors compared to diploid tumors (p <0.001). The presence of metastasis on presentation was significantly more frequent among biallelic and monoallelic tumors (p <0.001). Also, tumors greater in size than 4cm were significantly enriched for biallelic and monoallelic tumors (p = 0.002). No significant difference was observed for patient ethnicity, race and lymph node metastasis (Table 1).

**Table 1.** Clinicopathological data and associations

|  | Overall | Biallelic<br>Deletion | Monoallelic<br>Deletion | Low level<br>Gain | Diploid | P value |
| --- | --- | --- | --- | --- | --- | --- |
| No. pts | 440 | 17 | 116 | 7 | 300 | - |
| No. male (%) | 289 | 12 | 90 | 1 | 186 | <0.001 (Fisher's Exact Test) |
| Median Age (range) | 59 (26-90) | 64 (39-84) | 61.5 (34-90) | 62 (62) | 57.5 (26-88) | 0.047 (ANOVA) |
| No. ethnicity (%) |  |  |  |  |  | 0.557 (Fisher's Exact Test) |
| Hispanic or Latino | 24 | 0 | 7 (9) | 1 (16.7) | 16 (7.5) |  |
| Not Hispanic or Latino | 286 | 13 (100) | 71 (91) | 5 (83.3) | 197 (92.5) |  |
| Unknown | 130 | 0 | 39 | 1 | 88 |  |
| No. race (%) |  |  |  |  |  | 1.00 (Fisher's Exact Test) |
| Asian | 7 | 0 | 0 | 0 | 7 (2.4) |  |
| Black | 11 | 1 (7.1) | 1 (0.89) | 0 | 9 (3) |  |
| White | 416 | 13 (92.9) | 114 (99.1) | 7 (100) | 282 (94.6) |  |
| Unknown | 6 | 1 | 2 | 0 | 3 |  |
| No. AJCC Stage (%) |  |  |  |  |  | < 0.0001 (Kruskal-Wallis Test) |
| I | 200 (45.7) | 0 | 39 (33.9) | 2 (28.6) | 159 (52.8) |  |
| II | 44 (10) | 2 (13.3) | 7 (6.1) | 1 (14.3) | 34 (11.3) |  |
| III | 115 (26.3) | 5 (33.3) | 42 (36.5) | 3 (42.9) | 65 (21.6) |  |
| IV | 79 (18) | 8 (53.4) | 27 (23.5) | 1 (14.3) | 43 (14.3) |  |
| No. histological grade (%) |  |  |  |  |  | < 0.0001 (Kruskal-Wallis Test) |
| G1 | 7 (1.6) | 0 | 0 | 0 | 7 (2.3) |  |
| G2 | 181 (41.6) | 1 (6.7) | 35 (31) | 2 (28.6) | 143 (47.7) |  |
| G3 | 176 (40.5) | 5 (33.3) | 48 (42.5) | 3 (42.9) | 120 (40) |  |
| G4 | 71 (16.3) | 9 (60) | 30 (26.5) | 2 (28.6) | 30 (10) |  |
| GX | 2 | 0 | 1 | 0 | 1 |  |
| Presence of Mets on presentation (M1) | 77 | 8 | 28 | 0 | 41 | <0.001 (Fisher's Exact Test) |
| LN Mets (N1) | 16 | 3 | 5 | 0 | 8 | 1.00 (Fisher's Exact Test) |
| Size >4 cm | 325 | 15 | 98 | 5 | 207 | 0.002 (Fisher's Exact Test) |

### mRNA and Protein Expression levels of *CDKN2A/B*

The median *CDKN2A* and *CDKN2B* RNA expression levels were significantly lower in tumors with biallelic deletion compared to monoallelic and diploid tumors, however no significant difference was observed comparing monoallelic and diploid tumors (Fig. 2B). A similar trend was observed for *CDKN2A* protein expression data generated by RPPAs (Fig. 2A). Significantly lower protein expression levels were also observed among biallelic deleted tumors but not monoallelic tumors when compared to diploid tumors. Protein expression data for *CDKN2B* was not available.

**Figure 2.**
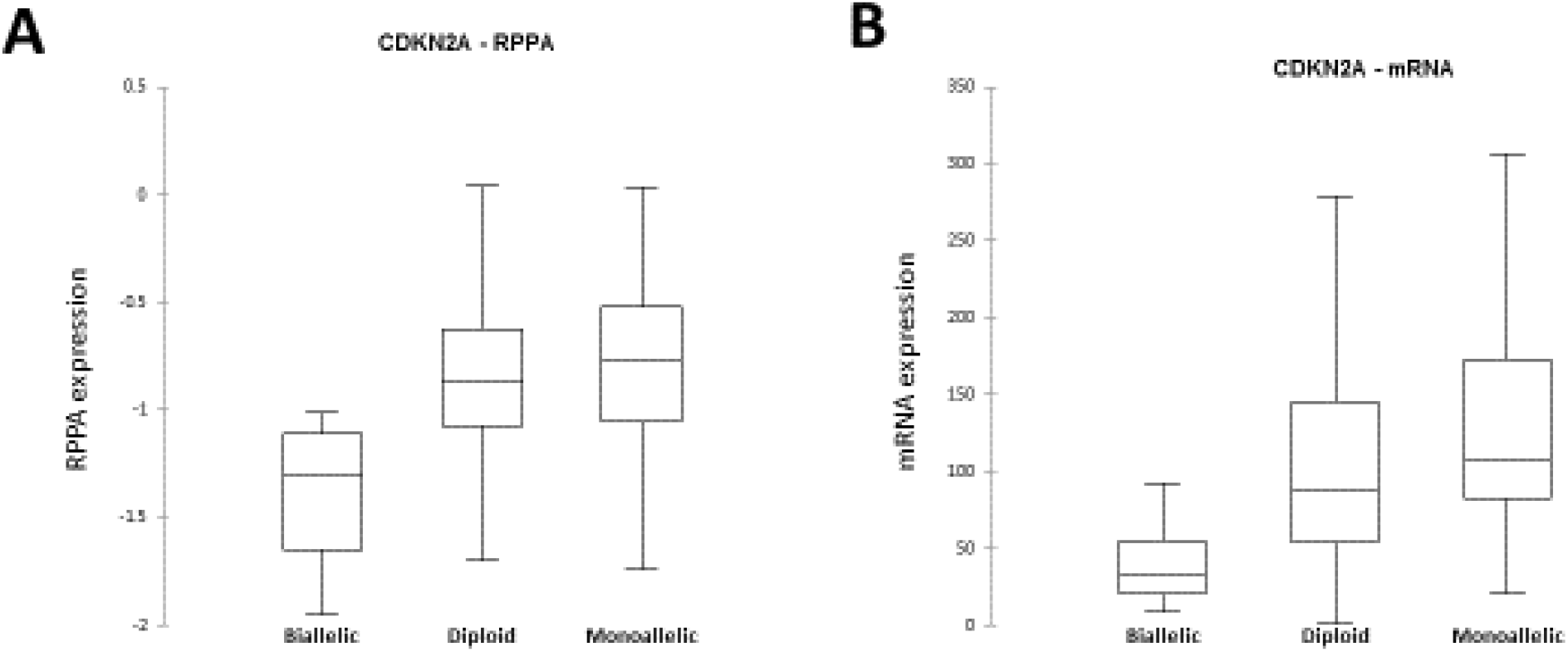
CDKN2A expression differences among CDKN2A/B biallelic deleted tumors versus monoallelic deleted and diploid tumors. Tumors with biallelic deletions of CDKN2A/B are associated with significant protein (A) and mRNA (B) underexpression of CDKN2A compared to diploid and monoallelic deleted tumors. No significant difference was observed between diploid and monoallelic deleted tumors at the protein and mRNA level.

### Prognostic Significance of Biallelic Inactivation of *CDKN2A/B*

Disease-free survival and cancer-specific survival was analyzed for association with *CDKN2A/B* status (See Materials and Methods). The median follow-up was 48.8 months (range 0.23-133.84) among survivors. Patients with *CDKN2A* biallelic deletion (median 30.45 months, 95% CI 17.96-40.59) had significantly worse cancer-specific survival than diploid tumors (median 45.11 months, 95% CI 43.50-50.95), however the difference in CSS of patients with monoallelic deletion was not significant (Figure 3A).

**Figure 3.**
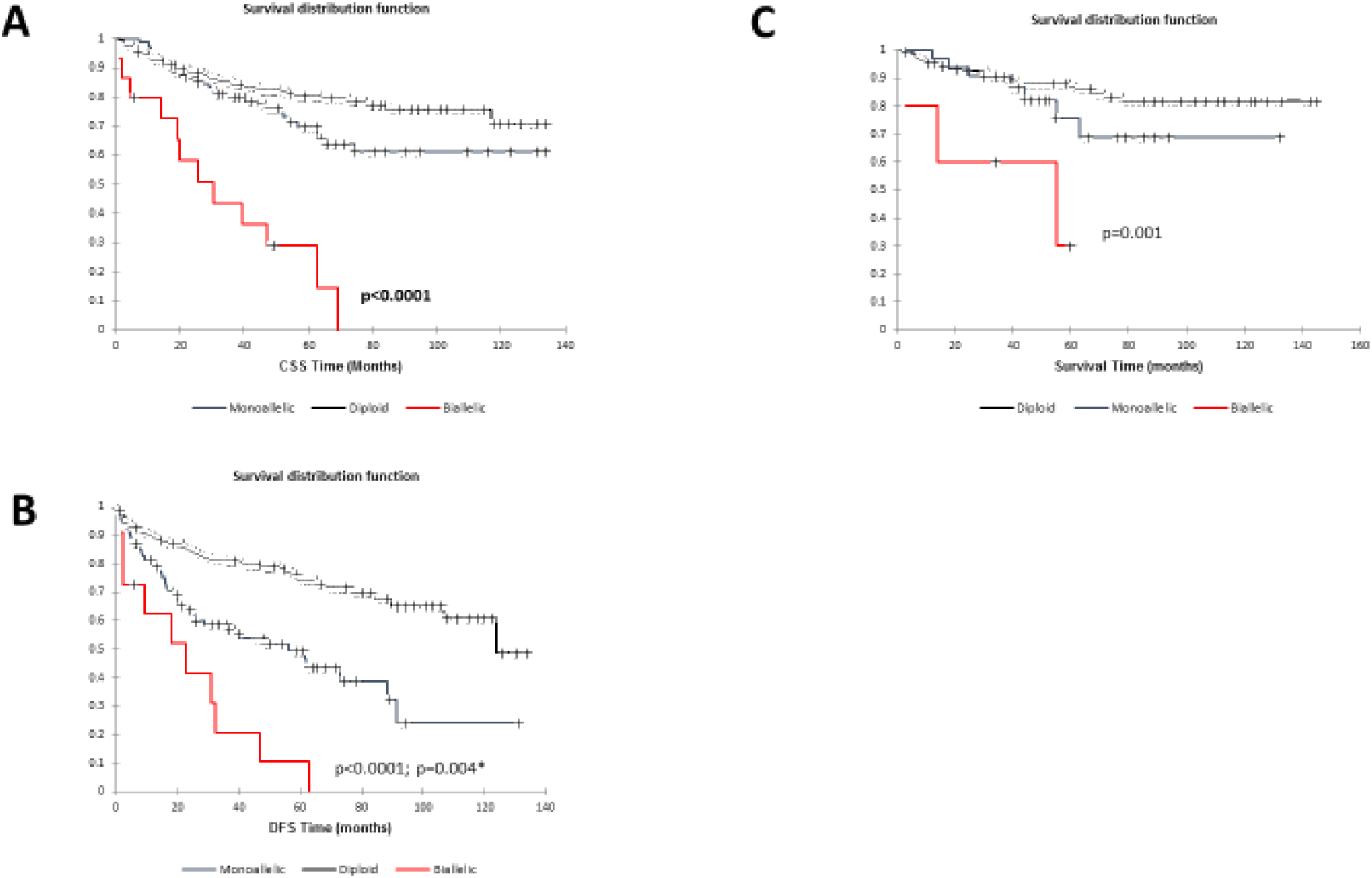
Biallelic and monoallelic deletions of CDKN2A/B significantly impact prognosis. A, patients with CDKN2A/B biallelic deletion had significantly worse cancer-specific survival than diploid tumors, however the difference in CSS of patients with monoallelic deletion was not significant. B, patients with CDKN2A/B biallelic and monoallelic deletion were significantly associated with worse DFS when compared to patients with diploid tumors. Biallelic deletion was also significantly associated with worse DFS when compared to patients with monoallelic deletion. C, Patients with biallelic deletions demonstrated significantly worse survival when compared to those with monoallelic deletions and diploid tumors. No significant difference was observed between monoallelic deleted tumors and diploid tumors.

Patients with *CDKN2A* biallelic (median 18.04 months, 95% CI 9.54-33.43) and monoallelic deletion (median 27.24 months, 95% CI 29.30-40.90) were significantly associated with worse DFS when compared to patients with diploid tumors (median 43.66 months, 95% CI 41.35-49.62; p<0.0001). Biallelic deletion was also significantly associated with worse DFS when compared to patients with monoallelic deletion (p=0.004; Figure 3B).

### Validation Set from Project GENIE and *Seto et al.*

Mutation of *CDKN2A* was observed in one case (n=237) in the Project GENIE dataset belonging exclusively to a metastatic tumor compared to its primary tumor. One *CDKN2B* mutation was observed in a separate primary tumor (Supplementary Table 1). Furthermore, in the Project GENIE dataset, one primary and 3 metastatic tumors (n=216) presented with *CDKN2A/B* biallelic deletion. Thirty-five (16.2%) tumors presented with monoallelic deletion of *CDKN2A/B* in the GENIE dataset.

The presence of biallelic deletions of *CDKN2A/B* in a third dataset of 240 ccRCCs was verified. A total of 5 cases (2.1%) were identified with biallelic deletions, followed by 34 cases (14.2%) with monoallelic deletions. Survival data was available for the 240 cases. Patients with biallelic deletions demonstrated significantly worse survival when compared to those with monoallelic deletions and diploid tumors (Figure 3C). No significant difference was observed between monoallelic deleted tumors and diploid tumors.

### Integrated *CDKN2A/B* Status and Whole-genome mRNA Profiling

To better elucidate the influence of *CDKN2A/B* allelic phenotypes on global mRNA expression, GSAA was conducted to assess the distribution of underexpressed and overexpressed sets of genes in relation to their genomic location and their copy number status with matched RNASeq expression data (see Materials and Methods). Three groups were compared: biallelic CDKN2A/B deleted tumors, monoallelic CDKN2A/B deleted tumors, and diploid tumors for CDKN2A/B. GSAA revealed 23 gene expression sets significantly enriched in association with biallelic CDKN2A/B tumors versus diploid tumors. Of these, 4 were associated with copy-number gains and 19 with monoallelic losses (p<0.05; q<0.25; Figure 1; Supplementary Tables S2 and S3). The most significantly altered gene set was loss of expression at 9p21, the region encoding *CDKN2A/B* (FDR = 0.0041), consisted with the allelic status of the subgroup being compared. This was followed by underexpressed gene expression on cytobands of chr9, chr4, chr6, chr13, chr14, and chr18. Overexpressed gene sets were represented on cytobands of chr7 and chr20. When comparing biallelic to monoallelic tumors, only two gene sets were enriched: 17q25 was significantly overexpressed and 4q22 was significantly underexpressed. Monoallelic tumors when compared to diploid tumors were enriched for overexpression of 21q22 and underexpression of chr9, chr4, chr6 and chr14 cytobands. 9p21 was also significantly underexpressed, however, ranked lower (7^th^ versus 1^st^) in its respective list of enriched regions.

### Integrated miRNA Profiling and *CDKN2A/B* Status

Genome-wide analysis of miRNAs for differential expression from the “Level 3” TCGA dataset comprising of 173 cases (See Materials and Methods) was conducted. Fifteen miRNAs (9 overexpressed and 6 under-expressed) were identified as significantly expressed in tumors with biallelic deletions of *CDKN2A/B* when compared to diploid tumors (Fig. 4). The well-known oncogenic miRNA, mir-21 was the most highly expressed in biallelic deleted tumors when compared to diploid tumors and when compared to monoallelic deleted tumors (Fig. 4B). mir-21 was previously demonstrated as a potential diagnostic and prognostic biomarker in ccRCC, with high levels associated with higher grade and stage and poor prognosis (33). This was followed by overexpression of members of the mir-183 miRNA family (mir-183 and mir-182), mir-9-1 and mir-9-2, mir-146b, mir-193a, and mir-625. The most underexpressed miRNA among biallelic deleted tumors was mir-126 (Fig. 4B), which was previously shown to be a marker of poor prognosis with relative low expression of mir-126 in metastatic ccRCC (34). This was followed by underexpression of mir-30c-2, mir-23b, mir-451, mir-204 and mir-144.

**Figure 4.**
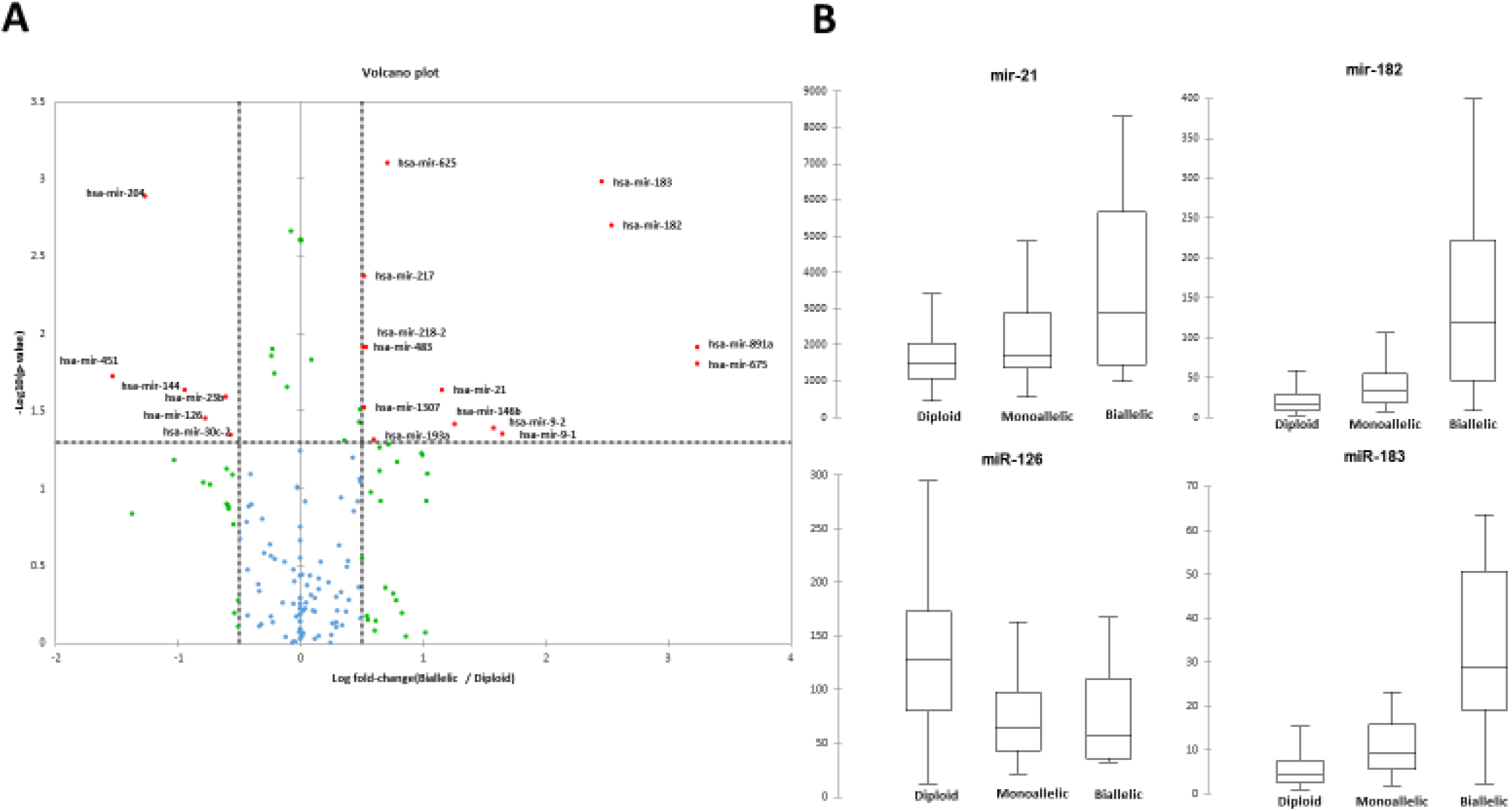
Differential expression of miRNAs in CDKN2A/B biallelic tumors versus monoallelic deleted and diploid tumors. A, volcano plot comparing miRNA expression between biallelic CDKN2A/B deleted tumors versus diploid tumors. B, boxplots of significantly expressed miRNAs miR-21 (overexpressed), mir-182 (overexpressed), mir-126 (underexpressed), mir-183 (overexpressed).

### Integrated Protein Profiling and Allelic Status of *CDKN2A/B*

“Level 3” RPPA data was queried through the cbio Cancer Genomics Portal for enrichment when comparing the three allelic phenotypes (See Materials and Methods). A list of 12 underexpressed and 10 overexpressed proteins were identified when comparing biallelic tumors to diploid tumors (Supplementary Table S4 and S5). Expectedly, CDKN2A protein expression was significantly underexpressed in biallelic tumors (Figure 2A), further confirming the impact of biallelic inactivation on the expression of *CDKN2A*. Among overexpression proteins were those involved in cell cycle processes (CCNB1, ASNS, YBX1), fatty acid synthesis (ACACA, FASN), iron transport (TFRC) and protein synthesis (EEF2, RPS6, RPS6KB1). Underexpression protreins were involved in metabolism (PRKAA1), cell growth (EGFR, ERBB3, ERRFI1), cell cycle (CDKN2A, PTPN11), vesicle trafficking (RAB11A, RAB11B), cell signaling (GAB2), tumor suppression (DPP4), and transcription (AR, STAT3).

### Integrated Pathway and gene signature analysis of *CDKN2A/B* Allelic Phenotypes

GSAA was also conducted to perform pathway analysis for enrichment of biological pathways, and to also cross-reference previously published gene signatures for significant overlap when comparing the three subgroups (See Materials and Methods). Biallelic deleted tumors were most significantly enriched for genes related to activation of ATR in response to replication stress (p=0.0067; Figure 5).

**Figure 5.**
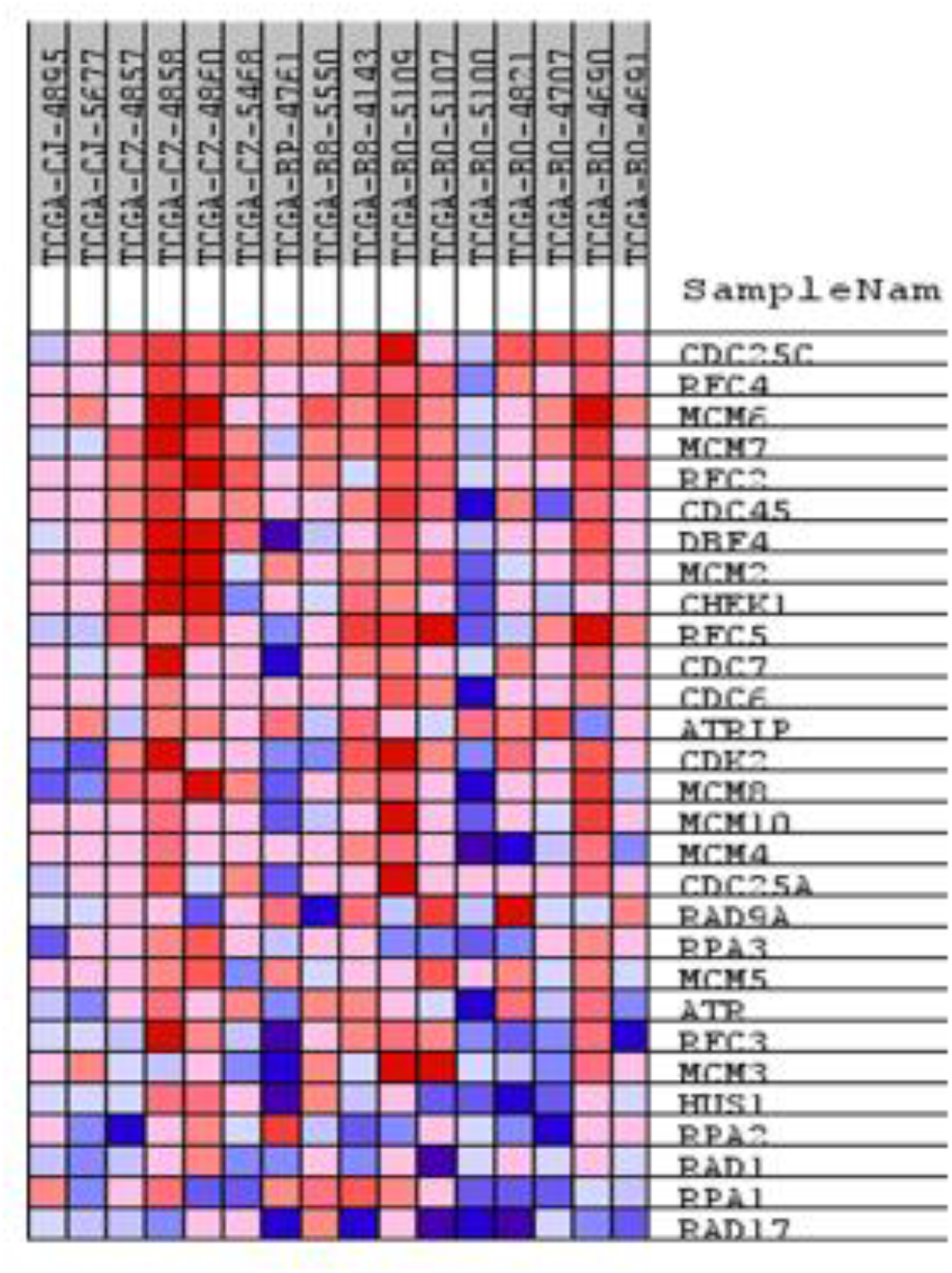
Heat map of genes associated with ATR in response to replication stress (red, overexpressed; blue underexpressed. Biallelic deleted tumors were most significantly enriched for genes related to activation of ATR in response to replication stress (p=0.0067)

The top and only significant miRNA signature at nominal p-value less than 0.01 identified when comparing both biallelic to monoallelic tumors and to diploid tumors was underexpression of miR-21 target genes (p=0.0073, p=0.0092, respectively). This observation is consistent with miR-21 presenting with the highest overexpression in biallelic tumors compared to both monoallelic and diploid tumors (Figure 4B).

## DISCUSSION

In this study, CDKN2A/B biallelic deleted tumors were characterized by assessing genome-wide copy number changes and integrated mutation data, gene, miRNA and protein expression, methylation and clinical data. A subset of tumors was identified with CDKN2A/B biallelic deletion in the TCGA dataset. A further set of tumors was identified to have monoallelic deletions of CDKN2A/B and the remaining cases were classified as either diploid or having a low-level gain of the locus (Table 1; Figure 1). Integrated mutation data identified a rare group of tumors with mutation of CDKN2A, but none with CDKN2B mutations.

Clinically, biallelic deleted tumors were associated with a higher median age and higher grade and stage. Furthermore, biallelic deleted tumors were significantly associate with presentation of metastasis (Table 1). To better define the clinical impact of biallelic and monoallelic deleted tumors, disease-free survival and cancer-specific survival was analyzed for association with *CDKN2A/B* status. Patients with *CDKN2A* biallelic deletion were associated with significantly worse cancer-specific survival than diploid tumors, while the difference in CSS of patients with monoallelic deletion was not significant (Fig. 3A). Additionally, *CDKN2A/B* biallelic and monoallelic were significantly associated with worse DFS when compared to patients with diploid tumors. Biallelic deletion was also significantly associated with worse DFS when compared to patients with monoallelic deletion (p=0.004; Fig. 3B). These results were further validated on an independent dataset (Fig. 3B).

To better elucidate the biological impact of CDKN2A/B biallelic deletion genome-wide mRNA expression data was integrated. Expression of genes in tumors with a CDKN2A/B biallelic deletion was compared to diploid tumors by GSAA. Twenty-three gene sets were identified as significantly altered between these 2 groups, providing rationale for a biologically distinct phenotype characterized by CDKN2A/B deletions.

Integration of miRNA expression data was conducted for further analysis. Fifteen miRNAs (9 overexpressed and 6 under-expressed) were identified as significantly expressed in tumors with mir-21 the most highly expressed in biallelic deleted tumors (Fig. 4B).

Further integration of protein (RPPA) data revealed 12 underexpressed and 10 overexpressed proteins when comparing biallelic tumors to diploid tumors (Supplementary Table S4 and S5).

## ACKNOWLEDGEMENTS

I would like to acknowledge the American Association for Cancer Research and its financial and material support in the development of the AACR Project GENIE registry, as well as members of the consortium for their commitment to data sharing. Interpretations are the responsibility of study author.

---

**Supplementary Table S1.**
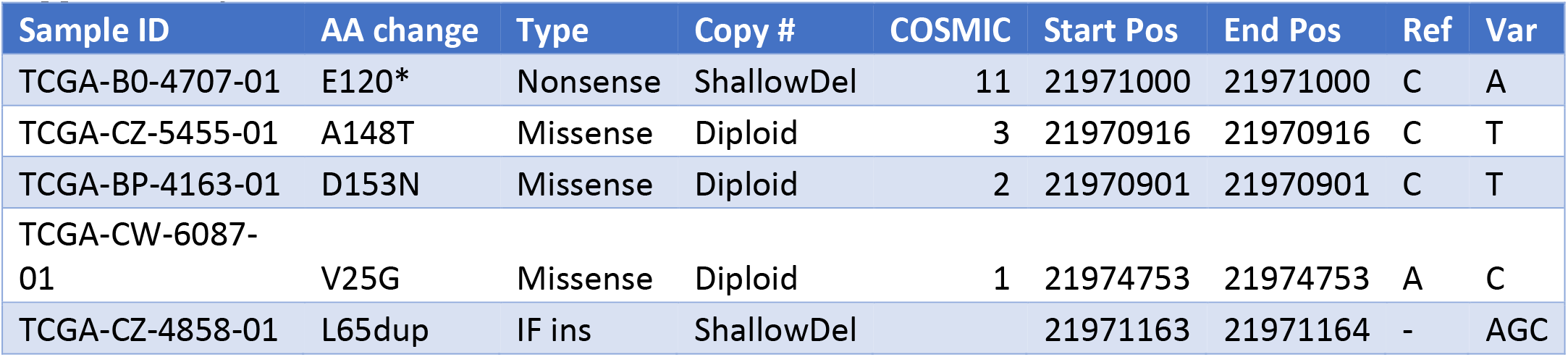
Mutations of CDKN2A in the ccRCC TCGA Dataset

| Sample ID | AA change | Type | Copy # | COSMIC | Start Pos | End Pos | Ref | Var |
| --- | --- | --- | --- | --- | --- | --- | --- | --- |
| TCGA-B0-4707-01 | E120* | Nonsense | ShallowDel | 11 | 21971000 | 21971000 | C | A |
| TCGA-CZ-5455-01 | A148T | Missense | Diploid | 3 | 21970916 | 21970916 | C | T |
| TCGA-BP-4163-01 | D153N | Missense | Diploid | 2 | 21970901 | 21970901 | C | T |
| TCGA-CW-6087-01 | V25G | Missense | Diploid | 1 | 21974753 | 21974753 | A | C |
| TCGA-CZ-4858-01 | L65dup | IF ins | ShallowDel |  | 21971163 | 21971164 | - | AGC |

**Supplementary Table S2.** GSAA gene sets significantly associated with overexpression in biallelic CDKN2A/B deleted tumors versus diploid tumors.

| NAME | SIZE | AS | NAS | NOM p-val | FDR q-val |
| --- | --- | --- | --- | --- | --- |
| CHR7P22 | 69 | 0.5971459 | 1.8540808 | 0.014150944 | 0.18766256 |
| CHR7Q22 | 109 | 0.5269004 | 1.8030012 | 0.034574468 | 0.23370832 |
| CHR20Q13 | 175 | 0.46380642 | 1.6888622 | 0.014749263 | 0.23834223 |
| CHR20Q11 | 109 | 0.5376253 | 1.855995 | 0.007853403 | 0.24594784 |

**Supplementary Table S3.** GSAA gene sets significantly associated with underexpression in biallelic CDKN2A/B deleted tumors versus diploid tumors.

| NAME | SIZE | AS | NAS | NOM p-val | FDR q-val |
| --- | --- | --- | --- | --- | --- |
| CHR9P21 | 30 | -0.73568153 | -2.1776624 | 0.001785714 | 0.004069265 |
| CHR9Q31 | 55 | -0.65324223 | -2.1379235 | 0 | 0.00560028 |
| CHR9Q21 | 49 | -0.66496634 | -2.0529897 | 0 | 0.013772152 |
| CHR4Q22 | 32 | -0.7035106 | -2.025251 | 0 | 0.01652819 |
| CHR9P22 | 35 | -0.63508874 | -2.0069199 | 0 | 0.017768454 |
| CHR9Q32 | 31 | -0.6608826 | -1.9855831 | 0.001834862 | 0.020762773 |
| CHR9Q22 | 83 | -0.5650191 | -1.9419571 | 0.00661157 | 0.026541194 |
| CHR9Q13 | 16 | -0.7019334 | -1.9196994 | 0.001602564 | 0.029769389 |
| CHR9P24 | 42 | -0.6143394 | -1.9421068 | 0 | 0.030332794 |
| CHR9P12 | 17 | -0.6130341 | -1.8355714 | 0.00189394 | 0.06804154 |
| CHR6Q15 | 16 | -0.70245415 | -1.8117217 | 0.007142857 | 0.078191064 |
| CHR4P12 | 20 | -0.67683864 | -1.7916638 | 0.00777605 | 0.08773007 |
| CHR13Q14 | 74 | -0.56234765 | -1.7565109 | 0.016638935 | 0.11506019 |
| CHR4Q21 | 80 | -0.5365679 | -1.6931602 | 0.026595745 | 0.1617375 |
| CHR13Q13 | 25 | -0.6533072 | -1.7084932 | 0.012027492 | 0.16268702 |
| CHR9P13 | 61 | -0.49950612 | -1.6998019 | 0.011994003 | 0.16290803 |
| CHR6P23 | 17 | -0.63557106 | -1.6341212 | 0.02443609 | 0.2043144 |
| CHR9Q12 | 16 | -0.60383695 | -1.6352379 | 0.016025642 | 0.21363144 |
| CHR18P11 | 63 | -0.51008016 | -1.6388583 | 0.02578269 | 0.2192122 |

**Supplementary Table S4.** Significantly underexpressed proteins in CDKN2A/B biallelic deleted tumors versus diploid tumors.

| Gene | Cytoband | difference | p-Value | q-Value |
| --- | --- | --- | --- | --- |
| PRKAA1_PT172 | 5p12 | -0.71 | 0.005484 | 0.0424 |
| EGFR_PY1068 | 7p12 | -0.59 | 0.003377 | 0.0355 |
| CDKN2A | 9p21 | -0.57 | 0.00007821 | 0.002366 |
| PTPN11_PY542 | 12q24 | -0.55 | 0.00006853 | 0.002366 |
| RAB11B | 19p13.2 | -0.54 | 0.000000171 | 0.00002071 |
| RAB11A | 15q22.31 | -0.54 | 0.000000171 | 0.00002071 |
| GAB2 | 11q14.1 | -0.49 | 0.0006643 | 0.0124 |
| ERBB3 | 12q13 | -0.45 | 0.0006331 | 0.0124 |
| DPP4 | 2q24.3 | -0.43 | 0.005611 | 0.0424 |
| AR | Xq12 | -0.39 | 0.004321 | 0.0387 |
| ERRFI1 | 1p36 | -0.34 | 0.000004401 | 0.000355 |
| STAT3_PY705 | 17q21.31 | -0.3 | 0.001911 | 0.024 |

**Supplementary Table S5.** Significantly overexpressed proteins in CDKN2A/B biallelic deleted tumors versus diploid tumors.

| Gene | Cytoband | difference | p-Value | q-Value |
| --- | --- | --- | --- | --- |
| YBX1 | 1p34 | 0.34 | 0.00002535 | 0.001534 |
| RB1_PS807_S811 | 13q14.2 | 0.41 | 0.001353 | 0.0193 |
| FASN | 17q25 | 0.44 | 0.0002103 | 0.005654 |
| YBX1_PS102 | 1p34 | 0.44 | 0.001986 | 0.024 |
| RPS6 | 9p21 | 0.45 | 0.001496 | 0.0201 |
| EEF2 | 19p13.3 | 0.5 | 0.006811 | 0.0485 |
| ASNS | 7q21.3 | 0.61 | 0.001353 | 0.0193 |
| ACACA | 17q21 | 0.61 | 0.005577 | 0.0424 |
| CCNB1 | 5q12 | 0.79 | 0.003102 | 0.0345 |
| TFRC | 3q29 | 1.22 | 0.000412 | 0.009971 |

## References

1. King SC, Pollack LA, Li J, King JB, Master VA. Continued Increase in Incidence of Renal Cell Carcinoma, Especially in Young Patients and High Grade Disease: United States 2001 to 2010. 2014;191:1665–70.

2. Chow W et al. Epidemiology and risk factors for kidney cancer. 2010;7:245–57.

3. Sun M, Thuret R, Abdollah F, Lughezzani G, Schmitges J, Tian Z, et al. Age-Adjusted Incidence, Mortality, and Survival Rates of Stage-Specific Renal Cell Carcinoma in North America: A Trend Analysis. 2011;59:135–41.

4. White NMA, Yousef GM. Translating Molecular Signatures of Renal Cell Carcinoma into Clinical Practice. J Urol. 2011;186:9–11.

5. Cancer T, Atlas G. Comprehensive molecular characterization of clear cell renal cell carcinoma. 24:3–9.

6. Hakimi AA, Ostrovnaya I, Reva B, Schultz N, Chen Y, Gonen M, et al. Adverse Outcomes in Clear Cell Renal Cell Carcinoma with Mutations of 3p21 Epigenetic Regulators BAP1 and SETD2: A Report by MSKCC and the KIRC TCGA Research Network.: 3259–67.

7. Lee HJ, Lee HY, Lee JH, Lee H, Kang G, Song JS, et al. Prognostic significance of biallelic loss of PTEN in clear cell renal cell carcinoma. J Urol [Internet]. Elsevier Ltd; 2014;192:940–6. Available from: http://www.ncbi.nlm.nih.gov/pubmed/24704020

8. Rayess H, Wang MB, Srivatsan ES. Cellular senescence and tumor suppressor gene p16. Int J Cancer. 2012;130:1715–25.

9. Harris SL, Levine AJ. The p53 pathway: positive and negative feedback loops. Oncogene; Oncogene. 2005;24:2899–908.

10. Zack TI, Schumacher SE, Carter SL, Cherniack AD, Saksena G, Tabak B, et al. Pan-cancer patterns of somatic copy number alteration. Nat Genet [Internet]. Nature Publishing Group; 2013;45:1134–40. Available from: /pmc/articles/PMC3966983/?report=abstract

11. Girgis AH, Iakovlev V V., Beheshti B, Bayani J, Squire J a., Bui A, et al. Multilevel whole-genome analysis reveals candidate biomarkers in clear cell renal cell carcinoma. Cancer Res. 2012;72:5273–84.

12. Beroukhim R, Brunet JP, Di Napoli A, Mertz KD, Seeley A, Pires MM, et al. Patterns of gene expression and copy-number alterations in von-Hippel Lindau disease-associated and sporadic clear cell carcinoma of the kidney. Cancer Res. 2009;69:4674–81.

13. Ismail E, Thomas K, Norman P, Stewart F, Ghulam N. Tumour suppressor gene (CDKNA2) status on chromosome 9p in resected renal tissue improves prognosis of localised kidney cancer. 2016;7.

14. El-Mokadem I, Lim A, Kidd T, Garret K, Pratt N, Batty D, et al. Microsatellite alteration and immunohistochemical expression profile of chromosome 9p21 in patients with sporadic renal cell carcinoma following surgical resection. BMC Cancer [Internet]. BMC Cancer; 2016;16:546. Available from: http://bmccancer.biomedcentral.com/articles/10.1186/s12885-016-2514-8

15. El-Mokadem I, Fitzpatrick J, Bondad J, Rauchhaus P, Cunningham J, Pratt N, et al. Chromosome 9p deletion in clear cell renal cell carcinoma predicts recurrence and survival following surgery. Br J Cancer [Internet]. Nature Publishing Group; 2014;111:1381–90. Available from: http://dx.doi.org/10.1038/bjc.2014.420

16. Jafri M, Wake NC, Ascher DB, Pires DE V, Gentle D, Morris MR, et al. Germline Mutations in the CDKN2B Tumor Suppressor Gene Predispose to Renal Cell Carcinoma. 2015;723–30.

17. Mermel CH, Schumacher SE, Hill B, Meyerson ML, Beroukhim R. GISTIC2. 0 facilitates sensitive and confident localization of the targets of focal somatic copy-number alteration in human cancers. 2011;1–14.

18. Li B, Dewey CN. RSEM: accurate transcript quantification from RNA-Seq data with or without a reference genome. 2011;

19. Zhao Q, Caballero OL, Davis ID, Jonasch E, Tamboli P, Yung WKA, et al. Tumor-Speci fi c Isoform Switch of the Fibroblast Growth Factor Receptor 2 Underlies the Mesenchymal and Malignant Phenotypes of Clear Cell Renal Cell Carcinomas. 2013;2:2460–72.

20. Escudier B, Camparo P, Doss DJ, Tannir NM. Next-Generation Sequencing of Translocation Renal Cell Carcinoma Reveals Novel RNA Splicing Partners and Frequent Mutations of Chromatin-Remodeling Genes. 3:4129–40.

21. Cancer T, Atlas G. Comprehensive molecular portraits of human breast tumours. 2012;1–10.

22. Manuscript A. NIH Public Access. 2014;6:1–34.

23. Cerami E, Gao J, Dogrusoz U, Gross BE, Sumer SO, Arman B, et al. In Focus The cBio Cancer Genomics Portal: An Open Platform for Exploring Multidimensional Cancer Genomics Data. 2012;

24. The AACR Project GENIE Consortium. AACR Project GENIE: Powering Precision Medicine Through An International Consortium, in preparation.

25. Sato Y, Yoshizato T, Shiraishi Y, Maekawa S, Okuno Y, Kamura T, et al. Integrated molecular analysis of clear-cell renal cell carcinoma. Nat Genet [Internet]. Nature Publishing Group; 2013;45:860–7. Available from: http://www.ncbi.nlm.nih.gov/pubmed/23797736

26. Xiong Q, Mukherjee S, Furey TS. GSAASeqSP: A Toolset for Gene Set Association Analysis of RNA-Seq Data. 2014;1–10.

27. Hamilton MP, Rajapakshe K, Hartig SM, Reva B, Mclellan MD, Kandoth C, et al. superfamily anchored by a central core seed motif. Nat Commun. Nature Publishing Group; 2013;4:1–13.

28. Ricketts CJ, Hill VK, Linehan WM. Tumor-Specific Hypermethylation of Epigenetic Biomarkers, Including SFRP1, Predicts for Poorer Survival in Patients from the TCGA Kidney Renal Clear Cell Carcinoma (KIRC) Project. 2014;9.

29. Carter H, Chen S, Isik L, Tyekucheva S, Velculescu VE, Kinzler KW, et al. Cancer-Specific High-Throughput Annotation of Somatic Mutations: Computational Prediction of Driver Missense Mutations. 2009;6660–7.

30. Reva B, Antipin Y, Sander C. Determinants of protein function revealed by combinatorial entropy optimization. 2007;8.

31. Reva B, Antipin Y, Sander C. Predicting the functional impact of protein mutations: application to cancer genomics. 2011;1–14.

32. Prmt M, Axis R, Marjon K, Cameron MJ, Quang P, Dorsch M, et al. Article MTAP Deletions in Cancer Create Vulnerability to Article MTAP Deletions in Cancer Create Vulnerability to Targeting of the MAT2A / PRMT5 / RIOK1 Axis. CellReports. The Authors; 2016;15:574–87.

33. Faragalla H, Youssef YM, Scorilas A, Khalil B, White NMA, Mejia-guerrero S, et al. The Clinical Utility of miR-21 as a Diagnostic and Prognostic Marker for Renal Cell Carcinoma. JMDI. Elsevier Inc.; 2012;14:385–92.

34. Khella HWZ, Scorilas A, Mozes R, Mirham L, Lianidou E, Krylov SN, et al. Low Expression of miR-126 Is a Prognostic Marker for Metastatic Clear Cell Renal Cell Carcinoma. Am J Pathol. American Society for Investigative Pathology; 2015;185:693–703.

